# Increased taxon sampling reveals thousands of hidden orthologs in flatworms

**DOI:** 10.1101/050724

**Authors:** José M. Martín-Durán, Joseph F. Ryan, Bruno C. Vellutini, Kevin Pang, Andreas Hejnol

**Author notes:** These authors contributed equally to this work (José M. Martín-Durán, Joseph F. Ryan). Corresponding author: Andreas Hejnol.

## Abstract

Gains and losses shape the gene complement of animal lineages and are a fundamental aspect of genomic evolution. Acquiring a comprehensive view of the evolution of gene repertoires is limited by the intrinsic limitations of common sequence similarity searches and available databases. Thus, a subset of the complement of an organism consists of hidden orthologs, those with no apparent homology with common sequenced animal lineages –mistakenly considered new genes– but actually representing rapidly evolving orthologs or undetected paralogs. Here, we describe Leapfrog, a simple automated BLAST pipeline that leverages increased taxon sampling to overcome long evolutionary distances and identify hidden orthologs in large transcriptomic databases. As a case study, we used 35 transcriptomes of 29 flatworm lineages to recover 3,427 hidden orthologs, some of them not identified by OrthoFinder, a common orthogroup inference algorithm. Unexpectedly, we do not observe a correlation between the number of hidden orthologs in a lineage and its ‘average’ evolutionary rate. Hidden orthologs do not show unusual sequence composition biases (e.g. GC content, average length, domain composition) that might account for systematic errors in sequence similarity searches. Instead, gene duplication and divergence of one paralog and weak positive selection appear to underlie hidden orthology in Platyhelminthes. By using Leapfrog, we identify key centrosome-related genes and homeodomain classes previously reported as absent in free-living flatworms, e.g. planarians. Altogether, our findings demonstrate that hidden orthologs comprise a significant proportion of the gene repertoire in flatworms, qualifying the impact of gene losses and gains in gene complement evolution.

## Introduction

Changes in gene complement are a fundamental aspect of organismal evolution (Ohno 1970; Olson 1999; Long, et al. 2003; De Robertis 2008). Current genome analyses estimate that novel genes, the so-called ‘taxonomically-restricted’ genes (TRGs) or ‘orphan’ genes –those without a clear homolog in other taxa– represent around 10–20% of the gene complement of most animal genomes (Khalturin, et al. 2009; Tautz and Domazet-Loso 2011). Although reported in some cases as non-functional open reading frames (ORFs) (Clamp, et al. 2007), TRGs are likely essential for the biology and evolution of an organism (Loppin, et al. 2005; Khalturin, et al. 2009; Knowles and McLysaght 2009; Li, et al. 2010; Colbourne, et al. 2011; Warnefors and Eyre-Walker 2011; Martin-Duran, et al. 2013; Palmieri, et al. 2014). The continuous increase in gene content is, however, balanced by a high rate of depletion among newly evolved genes (Tautz and Domazet-Loso 2011; Palmieri, et al. 2014) and by losses within the conserved, more ancient gene complement of animals (Kortschak, et al. 2003; Krylov, et al. 2003; Edvardsen, et al. 2005; Technau, et al. 2005).

Understanding the dynamic evolution of gene repertoires is often hampered by the difficulties of confidently identifying gene losses and gains. Gene annotation pipelines and large-scale comparisons (e.g. phylostratigraphy methods) largely rely on sequence-similarity approaches for gene orthology assignment (Alba and Castresana 2007; Domazet-Loso, et al. 2007; Tautz and Domazet-Loso 2011; Yandell and Ence 2012). These approaches depend on taxonomic coverage and the completeness of the gene databases used for comparisons. Although extremely useful in many contexts, sequence-similarity methods, such as Basic Local Alignment Search Tool (BLAST) (Altschul, et al. 1990), can be confounded in situations in which a gene evolves fast, is short, has an abundance of indels and/or exhibits similarity with other counterparts in only a small subset of residues (Moyers and Zhang 2015). These limitations, as well as intrinsic systematic errors of most algorithms (Liebeskind, et al. 2016), can generate significant biases when studying the evolution of protein-coding gene families (Elhaik, et al. 2006; Moyers and Zhang 2015, 2016). Accordingly, a proportion of the gene complement of an organism will be represented by genes that lack obvious affinity with homologs in the gene sets of the best annotated genomes –thus mistakenly considered potential TRGs– but actually representing fast evolving orthologs that we call hidden orthologs. This systematic error can potentially be overcome by more sensitive, although computationally intense, detection methods (e.g. profile HMMs, PSI-BLAST) (Altschul, et al. 1997; Eddy 2011; Kuchibhatla, et al. 2014), but also by increasing taxon sampling, which helps to bridge the long evolutionary gaps between hidden orthologs and their well-annotated, more conservative counterparts (fig. 1A).

**Figure 1.**
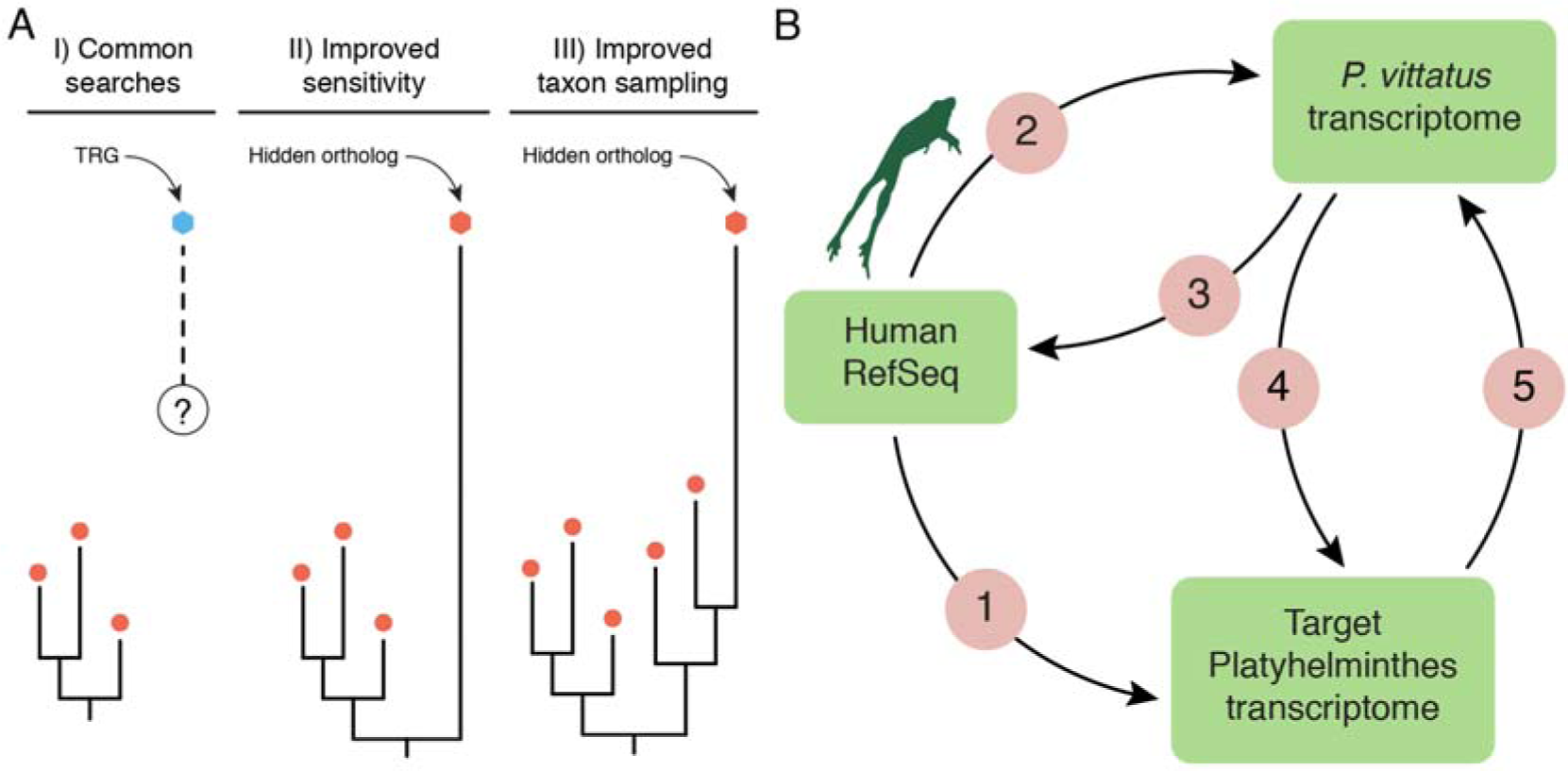
Hidden orthologs and the Leapfrog pipeline. (**A**) Taxonomically-restricted genes (TRGs) are genes with no clear orthology relationship (dashed line and question mark) to other known genes (e.g. orthology group of red dots). Improved sensitivity in the detection methods and/or improved taxon sampling can help uncover hidden orthology relationships, thus referring to these former TRGs as hidden orthologs. (**B**) The Leapfrog pipeline performs a series of reciprocal BLAST searches between an initial well-annotated dataset (e.g. human RefSeq), and a target and a ‘bridge’ transcriptomes. First, Leapfrog blasts the human RefSeq against the target (1) and the ‘bridge’ transcriptome (2), and identifies reciprocal best-hit orthologs between the human RefSeq and the ‘bridge’ (3). These annotated genes of the ‘bridge’ are then used to find orthologs in the target transcriptomes by reciprocal best BLAST hits (4 and 5). If these two pairs of reciprocal best BLAST hit searches are consistent between them, the gene in the target transcriptome is deemed a hidden ortholog.

Platyhelminthes (flatworms) is a morphological and ecologically diverse animal group characterized by significantly high rates of molecular evolution (Edgecombe, et al. 2011; Struck, et al. 2014; Laumer, Bekkouche, et al. 2015). Gene loss and orphan genes have been attributed to evolutionary changes leading to Platyhelminthes morphology (Berriman, et al. 2009; Martin-Duran and Romero 2011; Riddiford and Olson 2011; Tsai, et al. 2013; Breugelmans, et al. 2015). For instance, parasitic forms (e.g. tapeworms and flukes) have many unidentifiable genes and are reported to be missing myriad genes, including important developmental genes that are highly conserved in most other animals (Riddiford and Olson 2011; Tsai, et al. 2013). The presumed loss of critical genes has led to the inference that these animals have either developed alternative ways to implement critical steps in conserved pathways or that these pathways are no longer active (Wang, et al. 2011; Tsai, et al. 2013). A prime example is the loss of centrosomes in planarian flatworms, where the apparent absence of genes critical to the functioning of animal centrosomes was used as evidence supporting the secondary loss of these organelles in Platyhelminthes (Azimzadeh, et al. 2012).

Recently, two phylogenomic analyses have provided an extensive transcriptomic dataset for most platyhelminth lineages, in particular for those uncommon and less studied taxa that otherwise occupy key positions in the internal relationships of this group (Egger, et al. 2015; Laumer, Hejnol, et al. 2015). These important resources provide an ideal opportunity to address how increasing taxon sampling may improve the resolution of gene complement evolution in a fast evolving –and thus more prone to systematic error– animal group.

Here, we describe a BLAST pipeline, which we have called Leapfrog, that we have used to identify thousands of hidden orthologs across 27 different flatworms species by using a ‘slowly-evolving’ flatworm species as a ‘bridge’. Using this approach, we identify hundreds of hidden orthologs including tens of presumably lost centrosomal-related genes (Azimzadeh, et al. 2012) and several homeodomain classes previously reported as absent (Tsai, et al. 2013). Counter-intuitively, we show that the number of hidden orthologs does not correlate with the ‘average’ evolutionary rate of each particular species and unusual sequence composition biases, such as GC content, short transcript length and domain architecture that could affect BLAST searches. Instead, hidden orthologs appear to be generated by a combination of gene duplication events, weak positive selection and compensatory mutations between protein partners. Altogether, our findings demonstrate that a functionally relevant proportion of genes without clear homology are indeed hidden orthologs in flatworms, thus calling into question the current dogma of extensive gene loss within Platyhelminthes (Azimzadeh, et al. 2012; Tsai, et al. 2013). In a more general sense, our study provides an informative complementary approach for future high taxonomic resolution analyses of gene content in other phylogenetic clades.

## New Approaches

To identify hidden orthologs in large transcriptomic datasets we created Leapfrog, a simple pipeline that automates a series of BLAST-centric processes (fig. 1B). We started with a set of well-annotated sequences –a single-isoform version of the human RefSeq protein dataset– as queries and conducted a TBLASTN search of these sequences against each of our target flatworm transcriptomes (supplementary table 1, Supplementary Material online). Any queries that had zero BLAST hits with E-values less than our cutoff (0.01) were considered candidate hidden orthologs. We then looked for reciprocal best TBLASTX hits between these candidates and the transcriptome of the polyclad flatworm *Prostheceraeus vittatus*, a lineage that has evolved at a slower rate than most other flatworms in our dataset (as evidenced by root-to-tip branch lengths in (Laumer, Hejnol, et al. 2015)). If there was a reciprocal best BLAST hit in our ‘bridge’ transcriptome, the ‘bridge’ transcript was used as a query in a BLASTX search against the initial annotated human RefSeq protein dataset. If there was a human reciprocal hit, and the human sequence was the starting query, then we deemed the candidate a hidden ortholog.

## Results

### Leapfrog identified hundreds of hidden orthologs in flatworm transcriptomes

We assembled a dataset including 35 publicly available transcriptomes from 29 flatworm species, and incorporated the transcriptomes of the gastrotrich *Lepidodermella squamata*, the rotifer *Lepadella patella*, and the gnathostomulid *Austrognathia* sp. as closely related outgroup taxa. Under these conditions, Leapfrog identified a total of 3,427 putative hidden orthologs, 1,217 of which were unique and 636 were species-specific (fig. 2A, B; supplementary table 2, Supplementary Material online). In 30 cases (0.88% of the total recovered hidden orthologs), the same hidden ortholog and ‘bridge’ contig had a reciprocal best BLAST hit against two or more human proteins, likely paralogs/ohnologs (supplementary table 3, Supplementary Material online). From the annotation of their human ortholog, the flatworm hidden orthologs represented a wide array of different proteins, from genes involved in signaling transduction (e.g. GFRA3, a *GDNF family receptor alpha-3*) to DNA repair genes (e.g. BRCA2, the *breast cancer type 2 susceptibility protein*) and cytoskeleton regulators (e.g. COBLL1 or *cordon-bleu*). Indeed, alignments of recovered hidden orthologs with their human and *P. vittatus* counterparts show that many amino acid positions that differ between the human and the hidden ortholog products are conserved between *P. vittatus* and one or the other sequences (e.g., fig. 2C).

**Figure 2.**
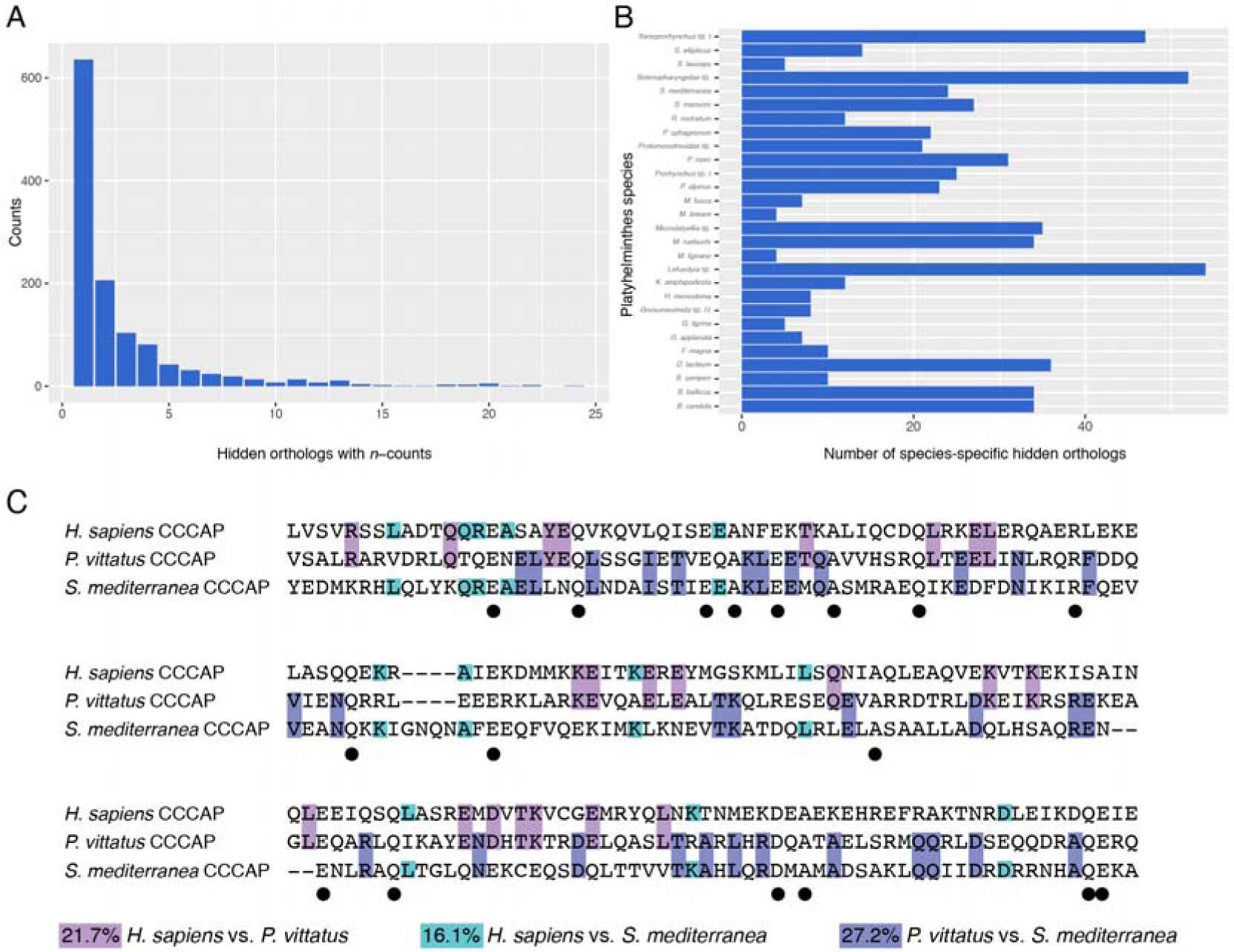
The Leapfrog pipeline recovers hundreds of hidden orthologs in Platyhelminthes. (**A**) Distribution of hidden orthologs according to their identification in one or more of the analyzed transcriptomes. Most of the hidden orthologs are unique of each lineage. (**B**) Distribution of species-specific hidden orthologs in each studied species. (**C**) Amino acid alignment of a fragment of the centrosomal protein CCCAP of *H. sapiens*, *P. vittatus* and *S. mediterranea*, and pairwise comparison of conserved residues. Positions that differ between the human and the hidden ortholog products are conserved between *P. vittatus* and one or the other sequences. Black dots indicate residues conserved among the three species.

In order to ensure that domain shuffling was not a source of false positives, we analyzed the domain content of 130 ‘bridge’ proteins and their corresponding human sequences from the *S. mediterranea* analysis (supplementary table 4, Supplementary Material online). In most cases (112; 86.15%), human and ‘bridge’ proteins had identical domain content. In the other 18 cases, InterProScan reported a domain in the ‘bridge’ ortholog that was not reported in the human protein or vice versa. Nonetheless, many of these may be due to InterProScan sensitivity. For example, in the case of BATF2, InterProScan identified the bZIP domain bZIP_1 (PF00170) in the human BATF2 and bZIP_2 (PF07716) in the ‘bridge’ sequence; however, bZIP_1 and bZIP2 are very similar and both domains are recovered in the identical region of BATF2 in hmmsearch results with very low E-Values of (3.2e-10 and 2.7e-06 respectively). In the other case, the LRRIQ1 (leucine rich repeats and IQ motif containing 1) gene, InterProScan identified only the IQ motif (PF00612) in the human gene, and only the LRR domains (PF13516, PF13855) in the ‘bridge’. In general, there was agreement between the domain architecture of the human and bridge sequences and we found no evidence suggesting that these results were inflated due to domain-shuffling events.

The number of hidden orthologs recovered in each particular lineage ranged from 41 in the rhabdocoel *Provortex sphagnorum* to 198 in the planarian *S. mediterranea* (fig. 3). The number of hidden orthologs varied considerably between different species belonging to the same group of flatworms. Within Tricladida, for instance, we identified 124 hidden orthologs in the marine species *Bdelloura candida*, 179 in the continenticolan species *Dendrocoelum lacteum* and 198 in the model species *S. mediterranea*. However, we only recovered 71 hidden orthologs for *Girardia tigrina*, a freshwater planarian related to *S. mediterranea*. We observed a similar issue in Macrostomorpha, Prorhynchida, and Rhabdocoela (fig. 3). Interestingly, the Leapfrog pipeline also reported hidden orthologs in the outgroup taxa (*Austrognathia* sp., 63; *L. patella*, 21; and *L. squamata*, 35) and *Microstomum lineare* (70), a flatworm lineage that shows a slower rate of evolutionary change than *P. vittatus* (Laumer, Hejnol, et al. 2015).

**Figure 3.**
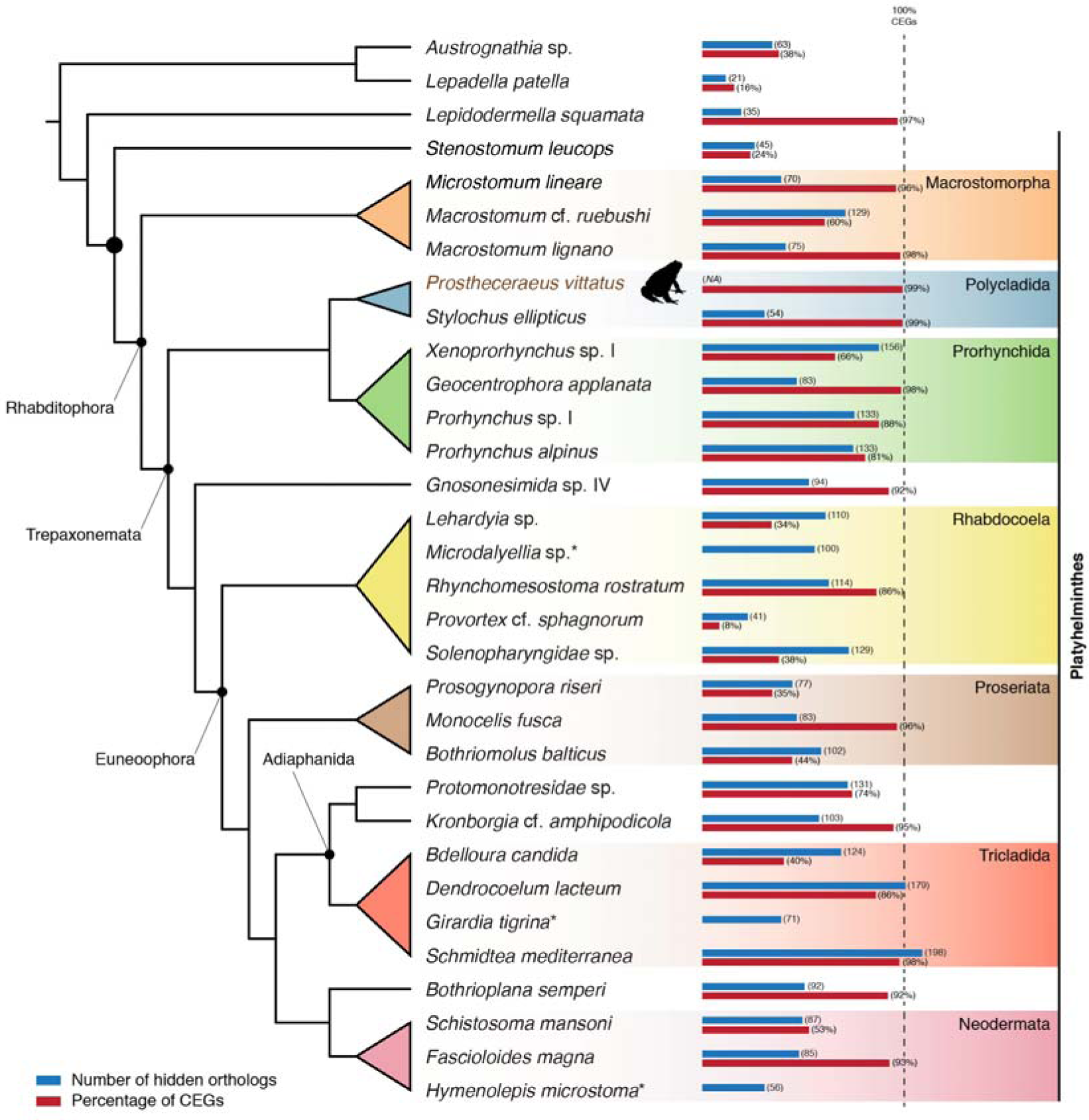
Distribution of hidden orthologs in the analyzed flatworm transcriptomes. The figure shows the total number of hidden orthologs in the analyzed transcriptomes in a phylogenetic context and with respect to their completeness (percentage of recovered core eukaryote genes, CEGs). The quality of the transcriptomes seems to be a limitation for the recovery of hidden orthologs in some flatworm lineages (e.g. *Provortex* cf. *sphagnorum*). However, the number of hidden orthologs is very species-specific.

To asses how the completeness of each transcriptome influenced Leapfrog results, we calculated the proportion of core eukaryotic genes (CEGs) (Parra, et al. 2007) present in each transcriptome. Consistent with the differences in sequencing depth (supplementary table 1, Supplementary Material online), we observed a broad range of CEG content between transcriptomes: from a reduced 8% in *P. sphagnorum* –the flatworm transcriptome with fewest recovered hidden orthologs– to an almost complete 99% of the polyclad *Stylochus ellipticus* and our ‘bridge’ species *P. vittatus* (fig. 3). Importantly, our dataset included highly complete transcriptomes (with > 85% CEGs) for each major flatworm group (Macrostomorpha, Polycladida, Prorhynchida, Rhabdocoela, Proseriata, Adiaphanida, Neodermata, Gnosonesimida and Bothrioplanida). The comparison of these highly complete transcriptomes with the other representatives of their respective groups showed that the number of recovered hidden orthologs was in many cases species-dependent. For instance, we recovered 83 putative hidden orthologs in *Geocentrophora applanata* and 133 in *Prorhynchus* sp. I, despite both prorhynchids having highly complete transcriptomes (fig. 3). The opposite case can be seen in the Macrostomorpha, where 70 (five species-specific) and 75 (four species-specific) hidden orthologs were recovered in *Microstomum lineare* and *Macrostomum lignano* respectively, both of which have highly complete transcriptomes. However, we identified 129 hidden orthologs (34 species-specific) in the closely related macrostomorph *Macrostomum* cf. *ruebushi*, whose transcriptome showed only a 60% of CEGs (fig. 3). These results together suggest that the number of hidden orthologs we recovered with Leapfrog is sensitive to the quality of the transcriptomes, but overall is variable even between closely related species.

We evaluated whether the use of a different ‘bridge’ transcriptome –with comparable completeness as *P. vittatus*– could be used to recover even more hidden orthologs in our datasets. We used the transcriptome of *M. lineare* because this species had the shortest branch in a published phylogenomic study (Laumer, Hejnol, et al. 2015). Using *M. lineare* as a ‘bridge’ we predicted hidden orthologs in the transcriptome of *S. mediterranea*, the lineage with the most hidden orthologs identified using *P. vittatus* as a ‘bridge’. Surprisingly, we only recovered 62 putative hidden orthologs under these conditions, as opposed to 198 when using *P. vittatus*, suggesting that evolutionary rate is not necessarily the best criteria for choosing a ‘bridge’ lineage. Noticeably, only 33 of the recovered 169 unique hidden orthologs overlapped between the two analyses, demonstrating the potential of using different transcriptomes as ‘bridges’ to identify additional hidden orthologs.

### Leapfrog identifies orthologs missed by OrthoFinder

In order to be certain that we were identifying orthologs that would be missed in a typical analysis, we compared our pipeline with the commonly deployed orthogroup inference algorithm OrthoFinder (Emms and Kelly 2015). To do this, we first performed an OrthoFinder analysis on human and the planarian *S. mediterranea*, and then evaluated the impact of adding the ‘bridge’ species *P. vittatus* to the calculation of orthogroups (fig. 3). Initially, OrthoFinder identified 5,638 orthogroups containing at least one sequence of *H. sapiens* and *S. mediterranea* (fig. 4A). The inclusion of the transdecoded ‘bridge’ transcripts of *P. vittatus* led to an increase in orthogroups that included both human and planarian sequences (5,816; fig. 4A). However, OrthoFinder only recovered 82.7% of the hidden orthologs identified by Leapfrog with a similar Evalue cutoff (fig. 4B). This is perhaps not surprising since OrthoFinder, as a general annotation tool, must balance sensitivity and specificity (Liebeskind, et al. 2016), while the Leapfrog pipeline is specifically designed to identify fast evolving orthologs.

**Figure 4.**
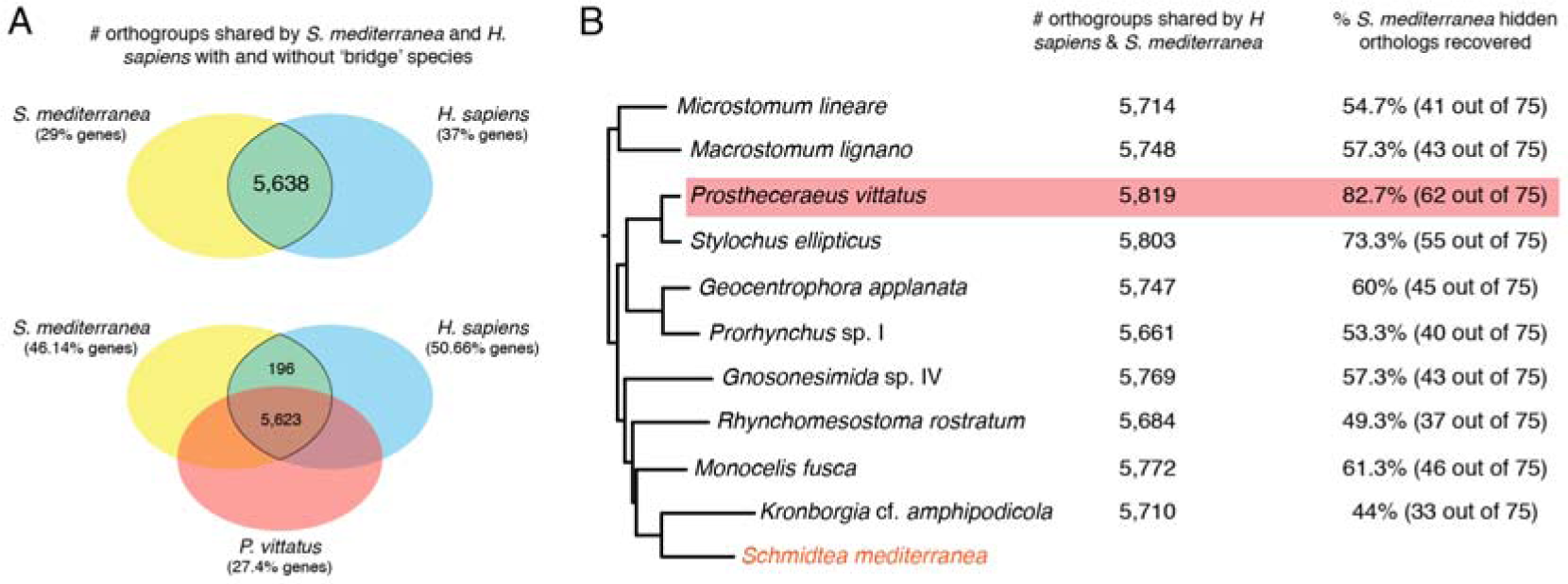
Comparison of Leapfrog with OrthoFinder. (**A**) Distribution of orthogroups identified by OrthoFinder when only *S. mediterranea* and human are considered (top) and when a ‘bridge’ transcriptome is added (bottom). The inclusion of an intermediate species increases the number of sequences assigned to orthogroups from a 29% to a 46.14% of the transcriptome, and the number of orthogroups between *S. mediterranea* and human from 5,638 to 5,819. (**B**) Effect of different ‘bridge’ transcriptomes in the identification of orthogroups and proportion of hidden orthologs recovered by Leapfrog present in orthogroups. As observed in Leapfrog, the orthology inference is sensitive to the ‘bridge’ transcriptome, but not strongly to the branch length of the ‘bridge’ species (depicted branch lengths are proportional to root-to-tip distances inferred from (Laumer, Hejnol, et al. 2015)). No single ‘bridge’ transcriptome included in an OrthoFinder analysis was able to recover all of the *S. mediterranea* hidden orthologs identified by the Leapfrog pipeline with similar E-value cutoffs.

To evaluate the impact of different ‘bridge’ species in OrthoFinder, we compared the number of hidden orthologs recovered by different bridge species. Bridge species were chosen only from deeply sequenced transcriptomes (with > 85% CEGs) and included a wide range of branch lengths (Laumer, Hejnol, et al. 2015). As observed with Leapfrog, the number of orthogroups greatly depended on the ‘bridge’ species and not so much on the branch length of the ‘bridge’ taxon: long-branch taxa recovered more orthogroups than shorter branches (e.g. *M. lignano* vs. *M. lineare*; *P. vittatus* vs. *M. lineare*; *Gnosonesimida* sp. IV vs. *R. rostratum*; *K.* cf *amphipodicola* vs. *Prorhyncus* sp. I). Importantly, none of the ‘bridge’ species were able to recover all the hidden orthologs that Leapfrog identifies in a *S. mediterranea* transcriptome (Brandl, et al. 2016) using a similar E-value cutoff (0.001) than OrthoFinder (Emms and Kelly 2015).

### The number of hidden orthologs does not relate to the branch length of each lineage

To investigate the parameters that might influence the evolutionary appearance and methodological identification of hidden orthologs in our dataset, we first performed a principal component analysis (PCA) including variables related to the quality and completeness of the transcriptome (number of sequenced bases, number of assembled contigs, mean contig length, and number of CEGs), the mean base composition of the transcriptome (GC content) and the evolutionary rate of each lineage (branch length, and number of identified hidden orthologs) (fig. 5A; supplementary table 5, Supplementary Material online). We observed that the first principal component (PC1) was strongly influenced by the quality of the transcriptome, while the second principal component (PC2) mostly estimated the balance between evolutionary change (branch lengths and hidden orthologs) and transcriptome complexity (GC content). The two first principal components explained 67% of the variance of the dataset, indicating that additional interactions between the variables exist (e.g. the GC content can affect sequencing performance (Dohm, et al. 2008; Benjamini and Speed 2012), and thus transcriptome quality and assembly).

**Figure 5.**
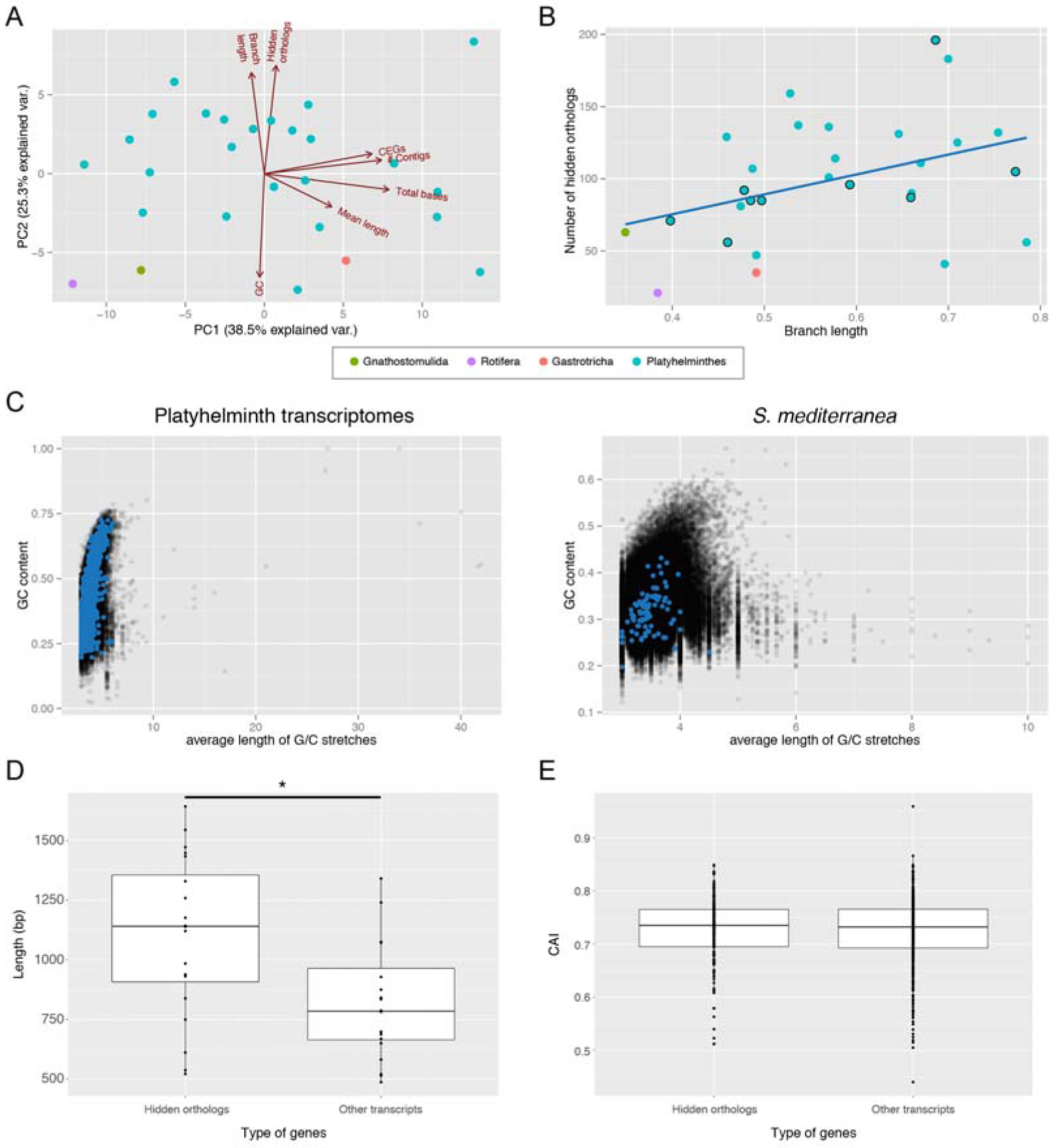
Hidden orthologs, evolutionary rates and sequence composition analyses. (**A**) Principal component analysis of the analyzed data showing the eigenvectors for each variable. The two first principal components (PC1, PC2) explain together 67.6% of the observed variability. (**B**) Number of hidden orthologs in relation to the branch length of each lineage (linear regression in blue; dots with external black line indicate the taxa with highly complete transcriptome). There is a low correlation between the two variables (R^2^=0.124). (**C**) GC content of each transcript plotted against its average length of G/C stretches considering all studied flatworm transcriptomes (left) and only *S. mediterranea* (right). The transcripts corresponding to hidden orthologs are in blue. Hidden orthologs do not differentiate from the majority of transcripts. (**D**) Average length of hidden orthologs compared to the average length of the other genes in transcriptomes with ≥ 85% CEGs. Hidden orthologs are significantly longer than the rest (Mann-Whitney test; p<0.05). (**E**) Codon Adaptation Index (CAI) of the hidden orthologs of the planarian species *B. candida*, *D. tigrina* and *S. mediterranea* compared with non-hidden orthologs. CAI index in hidden orthologs does not significantly differ from the rest of transcripts (Mann-Whitney test; p<0.05).

Despite the fact that the branch length of a given lineage and the number of putative hidden orthologs affected the dispersion of our data in a roughly similar manner, we did not detect a linear correlation (R^2^ = 0.131, p-value = 0.053; fig. 5B) between these two variables. When we considered those transcriptomes with similar completeness, the correlation between hidden orthologs and branch lengths was still weak (≥ 85% CEGs identified; R^2^ = 0.331, p-value = 0.04). This result supported our previous observation that lineages with similar branch lengths could exhibit remarkably different sets of hidden orthologs (fig. 3).

### Flatworm hidden orthologs do not show sequence composition biases

A recent report showed that very high GC content and long G/C stretches characterize genes mistakenly assigned as lost in bird genomes (Hron, et al. 2015). Given that GC content was implicated in our PCA, we tested whether high GC content and long G/C stretches were prevalent in our flatworm hidden orthologs. We first plotted the GC content and average length of the G/C stretches of all recovered hidden orthologs and compared them with all flatworm transcripts (fig. 5C). Hidden orthologs in flatworms do not show a significantly different GC content and average length of G/C stretches than the majority of transcripts. We confirmed this observation for each particular transcriptome of our dataset (fig. 5C; supplementary fig. 1, Supplementary Material online). These results indicate that hidden orthologs in flatworms are not characterized by the GC biases observed in birds (Hron, et al. 2015). Further investigation is needed to explore the possible causal relationship between hidden orthologs/branch lengths and GC content inferred from our PCA analysis (fig. 5A).

Systematic error in sequence-similarity searches is also associated with the length of the sequence and the presence of short conserved stretches (i.e. protein domains with only a reduced number of conserved residues). Short protein lengths decrease BLAST sensitivity (Moyers and Zhang 2015). We thus expected hidden orthologs to consist of significantly shorter proteins, as is seen in *Drosophila* orphan genes (Palmieri, et al. 2014). When analyzed together, the length of the flatworm hidden ortholog transcripts are not significantly different from that of the rest of the transcripts (supplementary table 6, Supplementary Material online). However, hidden orthologs are significantly longer than the rest of the transcripts when only high-coverage transcriptomes (≥ 85% CEGs identified) are considered (fig. 5D).

We next performed a domain-composition analysis of the 1,243 non-redundant human proteins homolog to the flatworm hidden orthologs (26 came from target contigs that matched several human proteins; supplementary table 2; Supplementary Material online), to address whether these sequences were enriched in particular sequence motifs that could hamper their identification by common sequence similarity searches. We recovered a total of 1,180 unique PFAM annotations, almost all of them present only in one (1,016) or two (112) of the identified hidden orthologs (supplementary table 7, Supplementary Material online). The most abundant PFAM domain (table 1) was the pleckstrin homology (PH) domain (PFAM ID: PF00169), which occurs in a wide range of proteins involved in intracellular signaling and cytoskeleton (Scheffzek and Welti 2012). PH domains were present in 11 of the candidate hidden orthologs. Most other abundant domains were related to protein interactions, such as the F-box-like domain (Kipreos and Pagano 2000), the forkhead-associated domain (Durocher and Jackson 2002), and the zinc-finger of C2H2 type (Iuchi 2001). These more abundant domains vary significantly in average length and number of generally conserved sites (table 1).

**Table 1.**
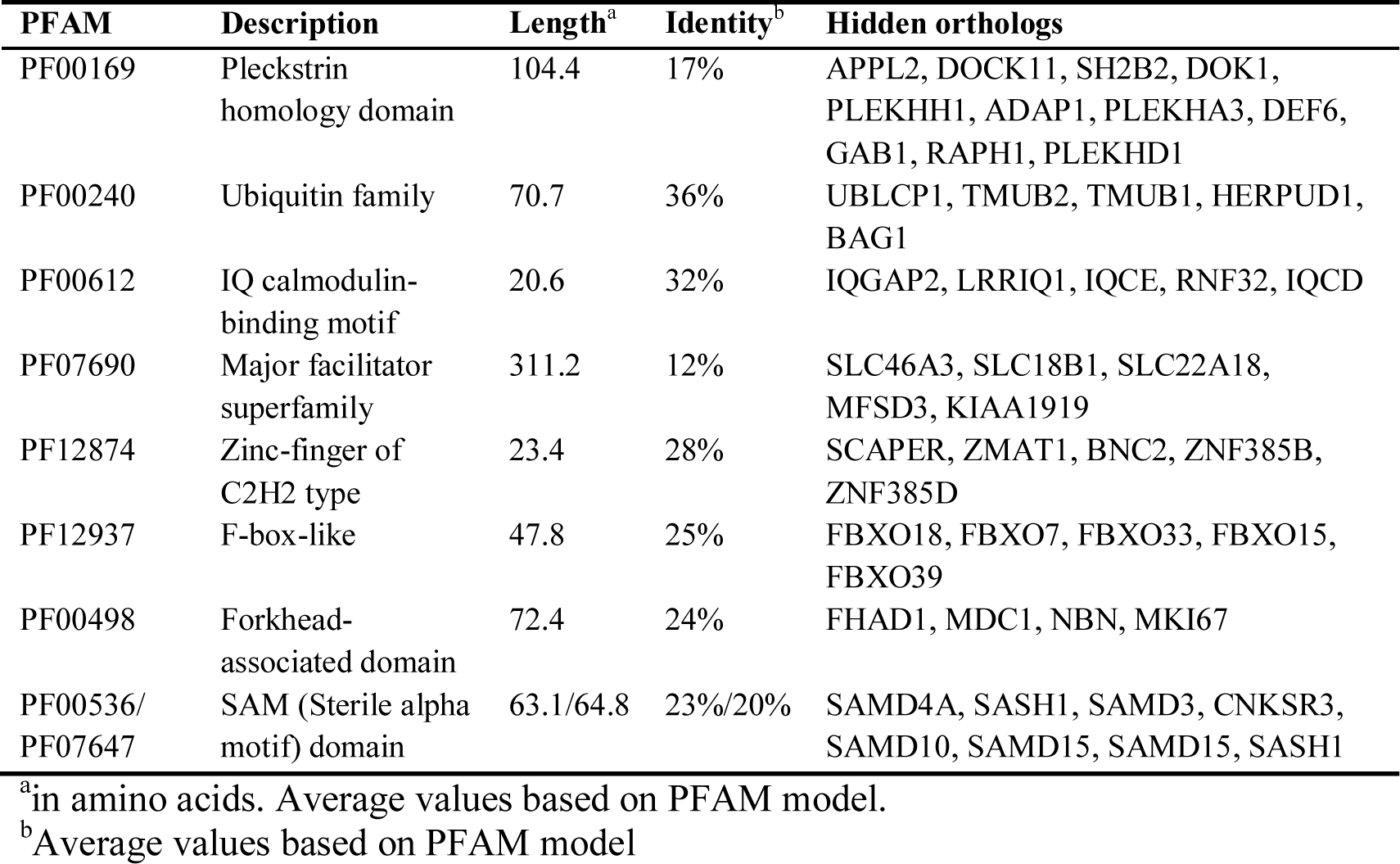
Most represented PFAM domains in flatworm hidden orthologs

Lastly, we looked to see if there were any patterns of codon usage associated with hidden orthologs. We did not observe a statistically significant difference between the codon adaptation index of hidden orthologs of the planarian species *B. candida*, *D. tigrina* and *S. mediterranea* and other open reading frames of these transcriptomes (fig. 5E). Altogether, these analyses indicate that hidden orthologs do not show intrinsic properties that could cause systematic errors during homology searches.

### The possible mechanisms driving hidden orthology in Platyhelminthes

To assess the contribution of duplication and divergence (Force, et al. 1999) towards the generation of hidden orthologs, we looked for paralogs of hidden orthologs in OrthoFinder orthogroups. Focusing on Tricladida (planarian flatworms), we counted the instances in which a hidden ortholog co-occurred in the same orthogroup with one or more sequences from the same species (fig. 6A). We observed that a hidden ortholog could indeed be interpreted as a fast-evolving paralog in 14–30% of the cases (depending on the species). For those one-to-one hidden orthologs of *S. mediterranea*, we calculated the number of non-synonymous substitutions per non-synonymous sites (*Ka*) and the number of synonymous substitutions per synonymous sites (*Ks*) in pairwise comparisons with their respective ortholog in the ‘bridge’ transcriptome (fig. 6B). Although for almost half of them the *Ks* value appeared to be saturated (*Ks* > 2), the *Ka*/*Ks* ratio for most of the rest was above or close to 0.5, which is often interpreted as sign of weak positive selection or relaxed constraints (Nachman 2006). Therefore, these findings suggest that duplication-and-divergence and weak positive selection may be forces contributing to hidden orthology in these data.

**Figure 6.**
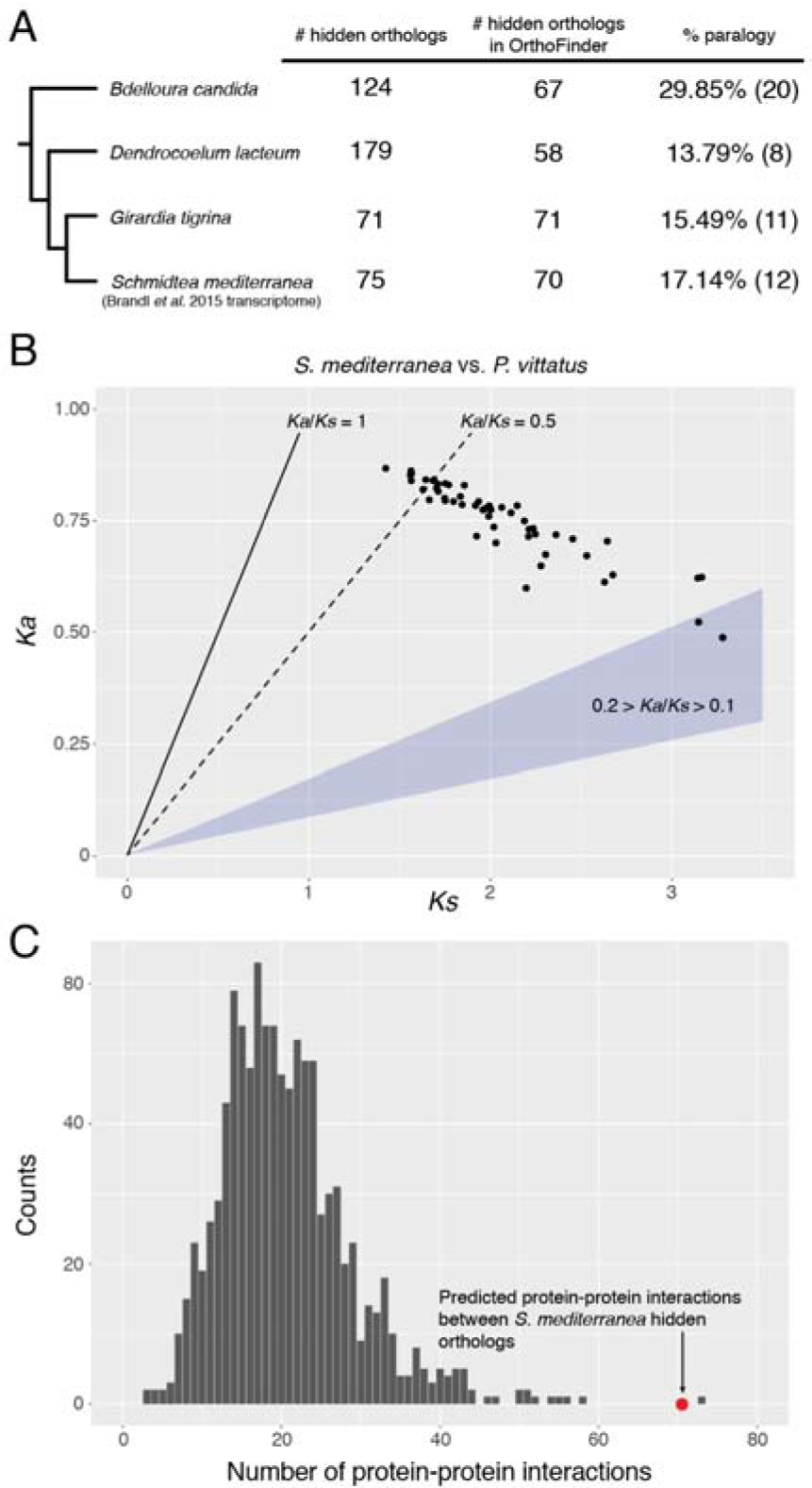
Level of paralogy and Ka/Ks values in triclad hidden orthologs. (**A**) Percentage of hidden orthologs identified by Leapfrog that are present in OrthoFinder and share orthogroup with other sequences of the same species. We dim these cases as probable fast evolving paralogs (hidden paralogy). (**B**) *Ka* and *Ks* values of 53 one-toone hidden orthologs of *S. mediterranea* compared with their respective homologues in the ‘bridge’ species *P. vittatus*. Although in almost half of these hidden orthologs the *Ks* value suggested saturation (*Ks* > 2), for most of the rest the *Ka*/*Ks* value was above or around 0.5 (dotted line), which can be a sign of weak positive selection or relaxed constraints. (**C**) Number of predicted protein-protein interactions in *S. mediterranea* hidden orthologs (red dot) compared with a distribution of interactions observed in 1,000 random samples of similar size (grey bars). Hidden orthologs show a significantly higher number of interactions, suggesting that complementary mutations between protein partners might drive hidden orthology in flatworms.

To test whether selective forces on particular cellular, molecular, or functional traits were responsible for hidden orthologs, we performed a gene ontology (GO) analysis of the non-redundant hidden orthologs identified in all flatworm transcriptomes. Amongst the wide spectrum of GO terms associated with hidden orthologs, binding and catalytic activities were the most abundant (supplementary fig. 2A–C, Supplementary Material online). We next performed a detailed GO analysis on the 198 hidden orthologs from the planarian *S. mediterranea* (supplementary fig. 2D–F, Supplementary Material online). The statistical comparison of the GO categories of the *S. mediterranea* hidden orthologs revealed 248 significantly (p < 0.05) enriched GO terms, 145 of them corresponding to the biological process category, 70 to the cellular component category, and 33 to the molecular function (table 2; supplementary table 8, Supplementary Material online). Interestingly, hidden orthologs were enriched for biological processes and cellular compartments related to mitochondrial protein translation and the mitochondrial ribosome respectively. Indeed, ribosomal proteins were amongst the most common hidden orthologs recovered from our dataset (supplementary table 2, Supplementary Material online). These findings suggest that mitochondrial genes show accelerated evolutionary rates (Solà, et al. 2015), which might be causing nuclear-encoded proteins that are exported to the mitochondrion adapt to this change (Barreto and Burton 2013).

**Table 2.** Enriched GO categories in *S. mediterranea* hidden orthologs

| **GO term** | **Description** | **E-value** |
| --- | --- | --- |
| **Biological process** |  |  |
| GO:0070124 | Mitochondrial translational initiation | 2.69E-12 |
| GO:0070126 | Mitochondrial translational termination | 4.07E-12 |
| GO:0070125 | Mitochondrial translational elongation | 6.05E-12 |
| GO:0032543 | Mitochondrial translation | 4.76E-07 |
| GO:0016064 | Immunoglobulin mediated immune response | 1.38E-06 |
| GO:0019724 | B cell mediated immunity | 1.38E-06 |
| **Molecular function** |  |  |
| GO:0001056 | RNA polymerase III activity | 9.55E-03 |
| GO:0005121 | Toll binding | 1.07E-02 |
| GO:0000989 | Transcription factor activity, transcription factor binding | 4.05E-02 |
| GO:0001635 | Calcitonin gene-related peptide receptor activity | 1.07E-02 |
| GO:0043237 | Laminin-1 binding | 1.07E-02 |
| GO:0005540 | Hyaluronic acid binding | 1.07E-02 |
| **Cellular compartment** |  |  |
| GO:0005761 | Mitochondrial ribosome | 3.87E-08 |
| GO:0000313 | Organellar ribosome | 6.08E-08 |
| GO:0005743 | Mitochondrial inner membrane | 9.09E-07 |
| GO:0019866 | Organelle inner membrane | 6.39E-06 |
| GO:0005762 | Mitochondrial large ribosomal subunit | 1.19E-05 |
| GO:0031966 | Mitochondrial membrane | 4.54E-05 |

To test if hidden orthologs are a result of rapid compensatory evolution, we investigated whether there were a disproportionate number of interactions between hidden orthologs in *S. mediterranea*. We used the BioGRID database (Chatr-Aryamontri, et al. 2015) to count the number of physical interactions (detected via experiments like affinity capture-luminescence, co-crystal structure, FRET—see BioGRID documentation for full list) between *S. mediterranea* hidden orthologs. We identified 71 such interactions in *S. mediterranea*, which is statistically significant when compared with 1000 random sets of human genes (one sample t-test; p-value: 2.2e-16; fig. 6C). These findings suggest that compensatory mutations in binding partners and/or otherwise interconnected proteins are contributing to the origin of hidden orthologs.

### The identified hidden orthologs fill out gaps in the flatworm gene complement

A previous study suggested the loss of an important proportion of centrosomal and cytoskeleton-related genes in the flatworms *M. lignano*, *S. mediterranea*, and *S. mansoni* (Azimzadeh, et al. 2012). We thus used an expanded Leapfrog strategy to identify possible hidden orthologs for that group of genes in our set of flatworm transcriptomes. First, we used a reciprocal best BLAST strategy to identify orthologs of the human centrosomal proteins in each of our transcriptomes under study, and thereafter we used Leapfrog to identify any hidden member of this original gene set. We recovered at least one reciprocal best BLAST hit for 56 of the 61 centrosomal genes, and identified fast-evolving putative orthologs in 19 of the 61 centrosomal genes (fig. 6). In total, the number of hidden orthologs identified was 58 (counting only once those for the same gene in the different analyzed *S. mediterranea* transcriptomes). Most importantly, we found the hidden orthologs for the genes CCCAP (SDCCAG8) and CEP192 in the planarian *S. mediterranea* (fig. 6; supplementary fig. 3 and supplementary fig. 4, Supplementary Material online), which were two of the five key essential centrosomal genes thought to be missing and essential for centrosome assembly and duplication (Azimzadeh, et al. 2012).

Hidden orthologs obtained in particular lineages could also be used as a “bridge” to manually identify their counterparts in other flatworm groups. For instance, we used the GFRA3 sequence from the fecampiid *Kronborgia* cf. *amphipodicola* and the FHAD1 (*fork head-associated phosphopeptide binding domain 1*) sequence from the rhabdocoel *Lehardyia* sp. to identify their putative orthologs in the planarian *S. mediterranea*. To explore the possibilities of this approach, we tried to manually identify in the planarian *S. mediterranea* classes of homeodomain genes previously reported as missing in free-living flaworms (Tsai, et al. 2013), using as a ‘bridge’ the orthologs found in the more conservative rhabditophoran species *M. lignano* and *P. vittatus*. We found orthologs for *gsc*, *dbx*, *vax*, *arx*, *drgx*, *vsx* and *cmp* in all these species (table 3; supplementary fig. 5 and supplementary fig. 6, Supplementary Material online), which places the loss of these homeodomain classes most likely at the base of the last-common neodermatan ancestor. Importantly, most of the classes absent in the transcriptomes of *P. vittatus* and *M. lignano* were also missing in *S. mediterranea*. The Hhex family was present in *P. vittatus*, but was not identified in *M. lignano* and *S. mediterranea*, and the Prrx and Shox families were present in *M. lignano*, but absent from *P. vittatus* and *S. mediterranea* transcriptomes. These observations suggest that many of the losses of homeobox genes occurred in the ancestors to the Rhabitophora and Neodermata, with only a few losses of specific gene classes in particular lineages of free-living flatworms.

**Table 3.** Presence/absence of hidden homeodomain genes in flatworms

| **Family** | ***M. lignano*** | ***P. vittatus*** | ***S. mediterranea*** |
| --- | --- | --- | --- |
| *Gsc* | — | Present | —<sup>1</sup> |
| *Pdx* | — | — | — |
| *Dbx* | Present | Present | Present |
| *Hhex* | — | Present | — |
| *Hlx* | — | — | — |
| *Noto* | — | — | — |
| *Ro* | — | — | — |
| *Vax* | Present | Present | Present |
| *Arx* | Present<sup>2</sup> | Present<sup>2</sup> | Present<sup>2</sup> |
| *Dmbx* | — | — | — |
| *Drgx* | Present<sup>2</sup> | Present<sup>2</sup> | Present<sup>2</sup> |
| *Prrx* | Present | — | — |
| *Shox* | Present | — | — |
| *Vsx* | Present | Present | Present (Kao, et al. 2013) |
| *Pou1* | — | — | — |
| *Cmp* | Present | Present | Present |
| *Tgif* | — | — | — |

<sup>1</sup>gene present in the sister species *S. polychroa* (Martin-Duran and Romero 2011)
<sup>2</sup>Orthology based on BBH, not well supported by phylogenetic relationships.

## Discussion

Our study reveals thousands of hidden orthologs in Platyhelminthes (fig. 2, 3), and thus illustrates the importance of a dense taxon sampling to confidently study gene losses and gains during gene complement evolution. Nevertheless, our approach is conservative and these results are likely an underestimation of the true number of hidden orthologs in these data.

Since our goal was to demonstrate how increased taxon sampling and the use of taxa with slow evolutionary rates can help identify fast evolving orthologs, we based our automated pipeline on BLAST searches (fig. 1B), by far the most common methodology for quickly identifying putative orthologs. However, other methods (e.g. profile HMM, PSI-BLAST) are more sensitive than BLAST when dealing with divergent sequences (Altschul and Koonin 1998; Eddy 1998), and have been shown, for instance, to recover homology relationships for many potential TRGs in viruses (Kuchibhatla, et al. 2014). Second, we based our identification of hidden orthologs on reciprocal best BLAST hits, a valid and widely used approach (Tatusov, et al. 1997; Overbeek, et al. 1999; Wolf and Koonin 2012), but with some limitations (Fulton, et al. 2006; Dalquen and Dessimoz 2013). Third, different ‘bridge’ transcriptomes generate different sets of hidden orthologs. This is an important observation, as it indicates that there might be natural circumstances (e.g., presence of hidden orthologs and missing genes), even in more conservatively evolving lineages, which contribute to the suitability of a particular transcriptome to act as a ‘bridge’. Therefore, an iterative approach in which all transcriptomes are used both as a ‘bridge’ and as a target will likely uncover even more hidden orthologs. Furthermore, we demonstrate that using hidden orthologs themselves as ‘bridge queries’ on other lineages can help recover even more new hidden orthologs (table 3). Finally, 16 out of the 35 analyzed transcriptomes contain less than 80% of core eukaryotic genes (fig. 3), and can be regarded as fairly incomplete (Parra, et al. 2009). All things considered, it is highly likely that the number of hidden orthologs in these flatworm lineages is far greater than what we are able to show in this study.

The recovered hidden orthologs have an immediate impact on our understanding of gene complement evolution in Platyhelminthes, and in particular on those lineages that are subject of intense research, such as the regenerative model *Schmidtea mediterranea* and parasitic flatworms (Berriman, et al. 2009; Wang, et al. 2011; Olson, et al. 2012; Sánchez Alvarado 2012). The identification of fast-evolving orthologs for the centrosomal proteins CEP192 and SDCCAG8 in *S. mediterranea* (fig. 6), as well as other core components in other flatworms lineages, indicates that the evolutionary events leading to the loss of centrosomes are probably more complex, or at least different from previously thought (Azimzadeh, et al. 2012). Similarly, the presence of presumably lost homeobox classes in *S. mediterranea* may affect our current view of gene loss and morphological evolution in flatworms (Tsai, et al. 2013). These two examples illustrate how our study and computational tools can serve the flatworm research community. The use of conservatively evolving flatworm lineages, such as *P. vittatus*, can improve the identification of candidate genes, as well as help with the annotation of the increasingly abundant flatworm RNAseq and genomic datasets (Berriman, et al. 2009; Wang, et al. 2011; Tsai, et al. 2013; Robb, et al. 2015; Wasik, et al. 2015; Brandl, et al. 2016). Therefore, we have now made available an assembled version of *P. vittatus* in PlanMine, an integrated web resource of transcriptomic data for planarian researchers (Brandl, et al. 2016). Importantly, the Leapfrog pipeline can also be exported to any set of transcriptomes/predicted proteins, and is freely available on GitHub (see Materials and Methods).

In our dataset, hidden orthologs are not significantly shorter, and do not exhibit either particular sequence composition biases (fig. 4) or protein domains (table 1) that could account for the difficulties in being detected by standard homology searches. Instead, hidden orthologs represent restricted fast evolving orthologs, which have been driven by either gene duplication events, weak positive selection and relaxed constraints, and/or by compensatory mutations between protein partners (fig. 6; supplementary fig. 3, Supplementary Material online). In some cases, hidden orthologs are associated with divergent biological features of Platyhelminthes (fig. 7; table 3). The fact that most of them are species-specific indicates that the gene complement of an organism is in fact heterogeneous, composed of genes evolving at different evolutionary rates (Wolfe 2004), sometimes much higher or much lower than the ‘average’ exhibited by that lineage.

**Figure 7.**
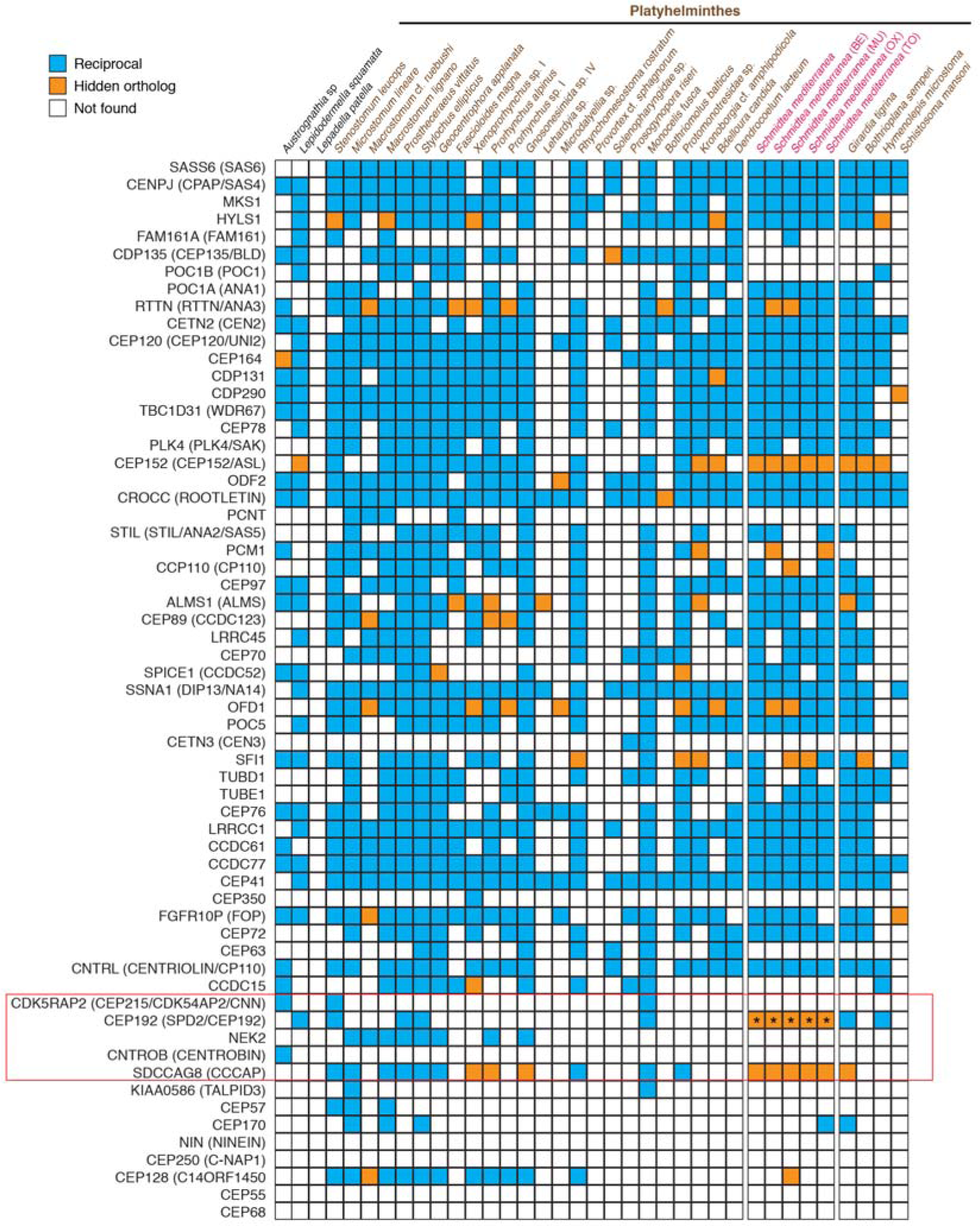
Hidden orthologs in the core set of centrosomal-related proteins. Presence (colored boxes) and absence (empty boxes) of the core set of centrosomal proteins (Azimzadeh, et al. 2012) in all the analyzed flatworm transcriptomes. Orthologs identified by direct reciprocal best BLAST hit are in blue boxes, and hidden orthologs are in orange. The CEP192 protein in the *S. mediterranea* transcriptomes (pink color code) is indicated by asterisks. These proteins were manually identified with the *G. tigrina* CEP192 as ‘bridge’ by reciprocal best BLAST hit. The five proteins essential for centrosomal replication are squared in red.

Previous studies suggested that more sensitive methods would reveal the real estimate of TRGs in animal genomes (Tautz and Domazet-Loso 2011). However, these methodologies are often time consuming and computationally intense, and thus hard to scale when dealing with large transcriptomes in a broad phylogenetic context. Our study proves that an alternative way to partially overcome this issue is by relying on improved taxon sampling, which is feasible as sequencing prices drop and the use of high-throughput sequencing becomes even more common in non-model organisms. Therefore, we envision a combination of both improved methodologies and expanded taxon sampling as the path to follow in future studies of gene complement evolution in animals.

Altogether, our study uncovers a so-far neglected fraction of the gene repertoire of flatworm genomes. Overlooked by common similarity searches, hidden orthologs include genes of biological relevance that were thought missing from the transcriptome/genome of most Platyhelminthes. These hidden genes are either maintaining ancestral functions despite very high mutation rates or are abandoning highly conserved ancestral functions but continuing to contribute to the biology of the organism. Either way, these results suggest that the prevalence of missing genes and orphan genes is likely exaggerated, and that caution is necessary in interpreting gene loss and gain when analyzing genomes.

## Materials and methods

### Macrostomum lignano *transcriptome*

Adult and juveniles of *M. lignano* were kept under laboratory conditions as described elsewhere (Rieger, et al. 1988). Animals starved for four days were homogenized and used as source material to isolate total RNA with the TRI Reagent (Life Technologies) following the manufacturer’s recommendations. A total of 1 μg was used for Illumina paired-end library preparation and sequencing in a HiSeq 2000 platform. Paired-end reads were assembled *de novo* with Trinity v.r20140717 using default settings (Grabherr, et al. 2011).

### Data set preparation

We downloaded the Human RefSeq FASTA file from the NCBI FTP site last updated on March 25, 2015 (ftp://ftp.ncbi.nlm.nih.gov/refseq/H_sapiens/H_sapiens/protein/protein.fa.gz). We also downloaded the gene2accession data file from NCBI, which was last updated on July 3, 2015 (ftp://ftp.ncbi.nlm.nih.gov/gene/DATA/gene2accession.gz). We then used the reduce_refseq script (available at https://github.com/josephryan/reduce_refseq) to generate a non-redundant Human RefSeq FASTA file with the following command: (reduce_refseq --fasta=protein.fa.gz --gene2accession=gene2accession.gz > HumRef2015.fa). This script prints only the first isoform for each Gene ID in the RefSeq FASTA file. The resulting file (available from the reduce_refseq repository) will be hereafter referred to as HumRef2015. Additionally, we downloaded the 28 RNA-Seq *de novo* assemblies from (Laumer, Hejnol, et al. 2015) and 6 additional *S. mediterranea* datasets from PlanMine v1.0 (Brandl, et al. 2016) on May 29, 2015. On July 14, 2015 we downloaded *Schistosoma mansoni, Hymenolepis microstoma,* and *Girardia tigrina* gene models from the Sanger FTP site. Further details on datasets are available in supplementary table 1 (supplementary Material online). All flatworm transcriptomes were constructed from adult tissue, except for *S. ellipticus*, *P. vittatus*, *G. applanata* and *Prorhynchus* sp. I, which included embryonic stages.

### Leapfrog Pipeline

All BLASTs were conducted using BLAST+ version 2.2.31 using multiple threads (from 2 to 10 per BLAST). We first ran a TBLASTN search using HumRef2015 as a query against the *Prostheceraeus vittatus* transcriptome (tblastn -query HumRef2015 -db Pvit -outfmt 6 -out Hs_v_Pv). We next ran a BLASTX search using the *Prostheceraeus vittatus* transcriptome as a query against the HumRef2015 dataset (blastx -query Pvit -db HumRef2015 -outfmt 6 -out Pv_v_Hs). We ran a series of TBLASTX searches using the *Prostheceraeus vittatus* transcriptome as a query against each of our target transcriptome database (e.g., tblastx -query “TRANSCRIPTOME” -db Pvit -outfmt 6 -out “TRANSCRIPTOME”_v_Pvit). Lastly, we ran a series of TBLASTX searches using our transcriptome databases as queries against the *Prostheceraeus vittatus* transcriptome (e.g., tblastx -query Pvit -db Sman -out Pvit_v_Sman -outfmt 6). The tab-delimited BLAST outputs generated above were used as input to the Leapfrog program (available from https://github.com/josephryan/leapfrog). The default E-Value cutoff (0.01) was used for all leapfrog runs. The leapfrog program identifies HumRef2015 proteins that fit the following criteria: (1) they have no hit to a target flatworm transcriptome, (2) they have a reciprocal best BLAST hit with a *Prostheceraeus vittatus* transcript, and (3) the *Prostheceraeus vittatus* transcript has a reciprocal best BLAST hit to the target flatworm transcriptome. The output includes the HumRef2015 Gene ID, the *Prostheceraeus vittatus* transcript and the target flatworm transcript. The annotation of the hidden orthologs is provided in supplementary table 2 (Supplementary Material online).

### OrthoFinder analyses

Single best candidate coding regions of the flatworm transcriptomes were predicted with TransDecoder v3.0.0 (Haas, et al. 2013). For *S. mediterranea*, we used the transcriptome sequenced by J. Rink’s lab at the MPI in Dresden (Brandl, et al. 2016). The resulting flatworm proteomes, together with HumRef2015, were used to identify orthologous groups with OrthoFinder v0.7.1 (Emms and Kelly 2015). Initially, an all-versus-all analysis was conducted. To define orthologous groups from a subset of transcriptomes, the clustering process was re-run with the –b option excluding the unwanted species.

### CEGMA analysis, transcriptome quality assessment, and statistics

Transcriptome completeness was evaluated with CEGMA (Parra, et al. 2007; Parra, et al. 2009). We could not run the CEGMA pipeline in the transcriptomes of *G. tigrina*, *Microdalyellia* sp. and *H. microstoma* due to an untraceable error. We calculated the contig metrics for each transcriptome assembly with TransRate (Smith-Unna, et al. 2015). Principal component analysis was performed in R and plotted using the ggplot2 package (Wickham 2009). For this analysis, branch lengths were the root-to-tip distances inferred from a maximum likelihood phylogeny previously reported (Laumer, Hejnol, et al. 2015).

### GC content, sequence length, CAI index, and Ka/Ks analyses

Custom-made scripts were used to calculate the GC content of hidden orthologs and transcripts of our dataset, the average length of the G/C stretches of each sequence, and the length of hidden orthologs and other transcripts. All scripts are available at https://github.com/josephryan/RyanLab/tree/master/2016-Martin-Duran_et_al. The codon usage matrices for *B. candida*, *D. tigrina* and *S. mediterranea* available at the Codon Usage Database (Nakamura, et al. 2000) were used as reference to calculate the ‘codon adaptation index’ with CAIcal server (Puigbo, et al. 2008). For each species, hidden orthologs were compared with three sets of transcripts generated by randomly choosing the same number of sequences than the number of hidden orthologs from the complete set of CDS sequences. To calculate *Ka* and *Ks* values, pairwise protein alignments of 53 one-to-one orthologs between *S. mediterranea* (hidden ortholog) and *P. vittatus* (‘bridge’ protein) were calculated with MAFFT v.5 (Katoh and Standley 2013). The software PAL2NAL (Suyama, et al. 2006) was used to infer codon alignments and trim gap regions and stop codons. *Ka* and *Ks* values were calculated with KaKs_calculator v2.0 (Wang, et al. 2010) with the default model averaging (MA) method. All values were plotted in R using the ggplot2 package (Wickham 2009).

### GO and InterPro analyses

GO analyses were performed with the human ortholog sequences from HumRef2015, using the free version of Blast2GO v3. Charts were done with a cutoff value of 30 GO nodes for the analyses of all hidden orthologs, and 10 GO nodes for the analyses of *S. mediterranea* hidden orthologs. Resulting charts were edited in Illustrator CS6 (Adobe). GO enrichment analysis of *S. mediterranea* hidden orthologs was performed with Blast2GO v3 comparing the GO annotations of the hidden orthologs against the GO annotations of the whole *S. mediterranea* transcriptome. InterProScan 5 was used to analyze the domain architecture of the recovered hidden orthologs using the human ortholog sequence, as well as 130 human and ‘bridge’ proteins to check for domain-shuffling events.

### Interaction analyses

We wrote a Perl script (biogrid.pl) to identify physical interactions in *S. mediterranea* hidden orthologs. The same script built 1,000 random gene sets (sampling genes from HumRef2015 that were current in Entrez Gene as of August 2016) and determined the number of physical interactions in each of these. The complete analysis can be repeated by running the biogrid.pl (https://github.com/josephryan/RyanLab/tree/master/2016-Martin-Duran_et_al), which produces R code that can be run to generate statistics reported herein. Results were plotted in R using the ggplot2 package (Wickham 2009).

### Multiple sequence alignments and orthology assignment

Full-length protein sequences of the human and *P. vittatus* SDCCAG8 gene were aligned to the SDCCAG8 cryptic ortholog recovered for *S. mediterranea*. Alignment was performed with MAFFT v.5 (Katoh and Standley 2013) using the G-INS-i option. Resulting alignment was trimmed between positions 319 and 494 of the human protein and edited with Illustrator CS6 (Adobe) to show the conserved residues between the three species. Multiple sequence protein alignments were constructed with MAFFT v.5 and spuriously aligned regions were removed with gblocks 3 (Talavera and Castresana 2007). Alignments are available at https://github.com/josephryan/RyanLab/tree/master/2016-Martin-Duran_et_al. Orthology assignments were performed with RAxML v8.2.6 (Stamatakis 2014) with the autoMRE option. The best fit models of protein evolution (CEP192: RtRev+I+G+F; CCCAP: JTT+G+F; Homeodomains: LG+G) were determined with ProtTest (Abascal, et al. 2005). Resulting trees were edited with FigTree and Illustrator CS6 (Adobe).

## Author’s contributions

JMMD and JFR designed the study. JMMD, AH, and KP collected material for the transcriptomes of *M. lignano, P. vittatus*, and *L. squammata*. JFR wrote the code of Leapfrog. JMMD, JFR and BCV performed the analyses. JFR, JMMD and AH wrote the manuscript. All authors read and approved the final manuscript.

## Acknowledgements

We thank the members of the Hejnol’s lab for support and discussions, and in particular Daniel Thiel and Anlaug Boddington for taking care of the *M. lignano* cultures. We appreciate the advance access given to platyhelmith transcriptomes by Gonzalo Giribet and Chris Laumer at the beginning of the project. We thank Axios Review (axiosreview.org) editor Titus Brown and two anonymous reviewers for the efficient and insightful reviewing of this paper. This research was funded by the Sars Centre core budget and the European Research Council Community’s Framework Program Horizon 2020 (2014–2020) ERC grant agreement 648861 to AH. JFR was supported by startup funds from the University of Florida DSP Research Strategic Initiatives #00114464 and the University of Florida Office of the Provost Programs, JMMD was supported by Marie Curie IEF 329024 fellowship.

